# Prognostic Significance of Potential Target Networks and P16INK4a In Early Diagnosis of Non-Small Cell Lung Cancer

**DOI:** 10.1101/2024.06.28.600894

**Authors:** Varnika Kundadka, Radha Vatsa, Navrinder Kaur

**Affiliations:** People’s Laboratory of Bioinformatics and Systems Biology, Quick IsCool, Aitele Research LLP, Bihar-843302, India; Department of Medicine, Imperial College London; Department of Bioinformatics, Patkar Varde College, University of Mumbai

**Keywords:** NSCLC, Biomarker, Bioinformatics, Lung cancer, P16, diagnostic markers, therapeutic biomarkers

## Abstract

This study aimed to identify the prognostic signature of p16INK4a and related genes in non-small cell lung cancer (NSCLC). Since non-small cell lung cancer shows a poor prognosis, the 5-year survival rate in advanced stages is low. This emphasizes the need to identify more reliable diagnostic biomarkers for the early diagnosis of NSCLC and understand the molecular mechanisms underlying the progression of the disease. The role of P16 in essential cellular pathways are still unknown. Understanding its function and the mechanisms underlying its dysregulation could potentially lead to the development of targeted therapies for certain cancers. To address this, bioinformatics analyses revealed differential expression of genes in lung tumour tissues vs control samples. The interactions among the identified DEGs can help better understand the underlying molecular mechanism of NSCLC. We investigated the correlation between p16INK4a expression and genes involved in the relevant interactive pathways in early-stage NSCLC. These can serve as potential diagnostic and therapeutic biomarkers for NSCLC.

## Introduction

Lung cancer is a formidable global health issue, representing one of the most common and deadly forms of cancer. According to global cancer statistics (2020) by WHO, lung cancer accounts for 11.4% of all cancers as shown in figure 1 (*Cancer Today*, n.d.). In men, it ranks the highest with 14.3% of all reported cases and third amongst women with 8.4% of the reported cases worldwide. Lung cancer also has the highest mortality rate of 18% among all cancers. GLOBOCAN India statistics suggest that lung cancer is rated sixth highest with approximately 72,000 new cases reported in 2020 as shown in out of which 66,000 deaths are reported indicating higher mortality rates. Therefore, it is necessary to study the early diagnostic and treatment methods to address the alarming rate of increase in the affected population.

**Figure 1.**
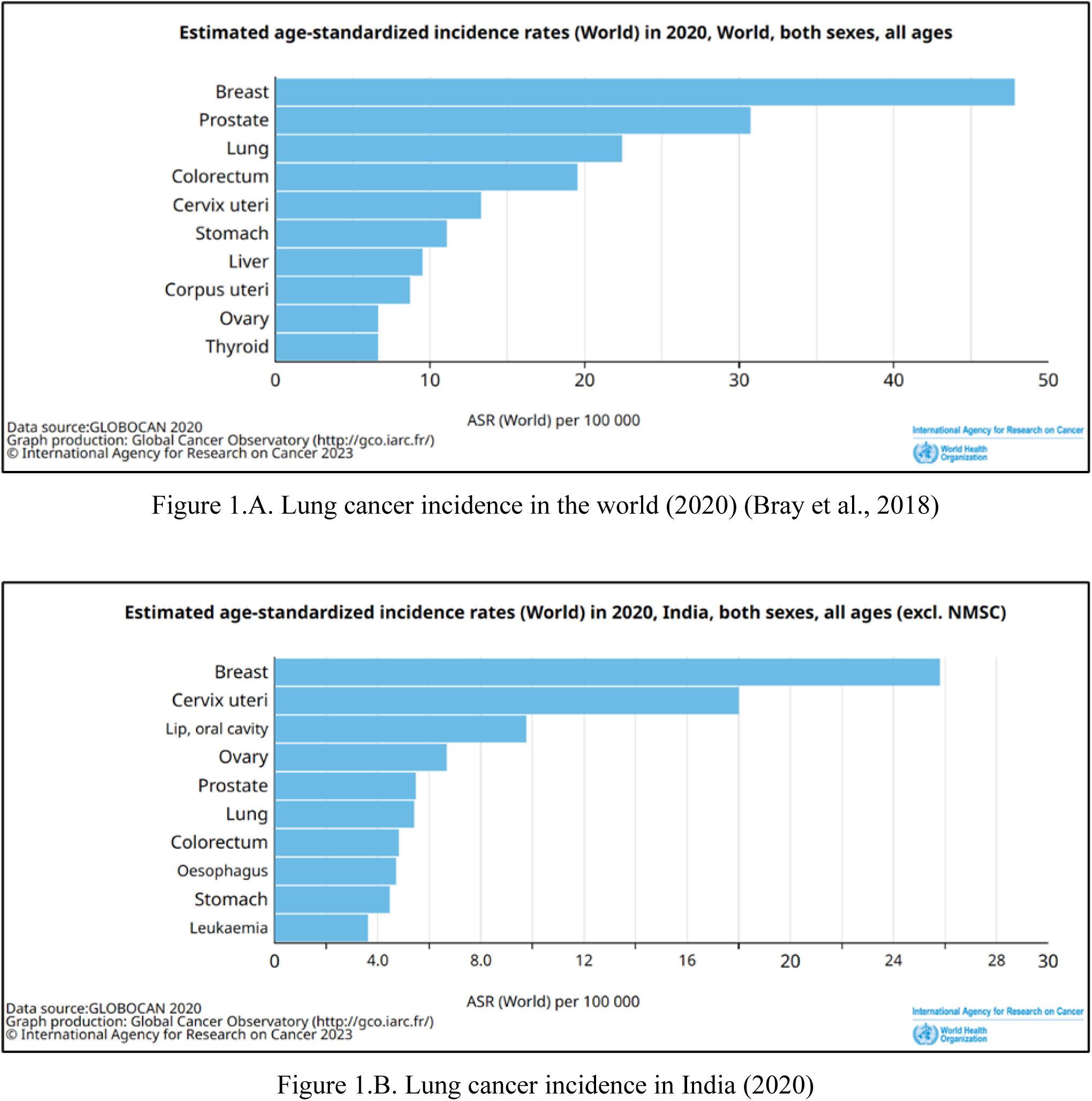
Estimated incidence rates in 2020. Bar graphs derived from International Agency for Research on Cancer (IARC) WHO. X axis marks the age standardized mortality rates. The Y-axis represents the type of cancer in descending order of ASR. The data includes all sexes at all ages. Figure 1.A. is the bar graph for lung cancer incidence in the world (2020) Figure 1.B. is the bar graph for lung cancer incidence in India (2020).

There are two primary types of lung cancer: Non-Small Cell Lung Cancer (NSCLC) and Small Cell Lung Cancer (SCLC). NSCLC is the most prevalent, comprising approximately 80% of all lung cancer cases (Garinet et al., 2022). NSCLC encompasses various histological subtypes, with adenocarcinoma, squamous cell carcinoma, and large cell carcinoma being the primary ones (Liu et al., 2019). Adenocarcinoma has shown an increasing trend in recent years and is now the most common subtype. While smoking remains the primary risk factor, environmental exposure, genetic predisposition, and certain lung diseases also play a significant role in its development.

The prognosis of non-small cell lung cancer (NSCLC) is still not optimistic. NSCLC patients bear poor prognosis, as the 5-year survival rate is 62.8%, for patients with stage I disease and is decreased to 8.2% for those with stage IV disease (*Lung and Bronchus Cancer — Cancer Stat Facts*, n.d.). In India, Noronha et al., 2012 indicate that 14% of their surveyed cancer patients were incorrectly diagnosed while 43% were already at Stage III during diagnosis which dramatically influences the success of the treatment increasing mortality rates. This emphasizes the need to identify more reliable diagnostic biomarkers for the early diagnosis of NSCLC and understand the molecular mechanisms underlying the progression of the disease.

The past decade has witnessed the discovery of multiple molecular aberrations that drive lung cancer growth. Tumour suppressors are genes that assist in regulating cellular proliferation and inhibiting the onset of malignancy. They fulfil a pivotal function in upholding the cellular life cycle’s stability and restraining unrestrained cellular partitioning. Deactivation of these genes consequently results in the formation of tumours. One such tumour suppressor gene that has been heavily associated with NSCLC is P16INK4a (Tuo et al., 2018). The p16INK4a/p16 gene produces a protein named cyclin-dependent kinase inhibitor 2A (CDKN2A) (Jiao et al., 2018). This protein acts as a suppressor of the cell cycle, particularly controlling the G1 stage of the cell cycle. CDKN2A inhibits cyclin-dependent kinase [CDK4 and CDK6] with D-type cyclins and its long-term expression leads to cell senescence (Asghar et al., 2015). It hinders the phosphorylation of the retinoblastoma (Rb) protein, which results in the halt of the cell cycle and the prevention of progression from the G1 to the S stage, preventing uncontrolled cell growth and division (Antonucci et al., 2014). Lung cancer risk factors have been shown to increase the likelihood of p16INK4a alterations (Li et al., 2020). Loss of function of p16INK4a relieves the inhibitory constraints on CDK4 and CDK6, thus facilitating continuous phosphorylation of Rb (Dick et al., 2018). This leads to deactivation of Rb’s growth-suppressive functions. Due to its critical role in cell cycle regulation and cancer development, the p16INK4a gene has become a subject of significant interest in cancer research. With the above-mentioned developments, the role of P16 in essential cellular pathways are still unknown. Understanding its function and the mechanisms underlying its dysregulation could potentially lead to the development of targeted therapies for certain cancers.

Bioinformatics tools are utilized to identify the differential expressed genes (DEG) (Gilbert, 2000). By comparing gene expression patterns in lung cancer tissue samples to normal lung tissue, researchers can pinpoint the genes that are overexpressed or under expressed in NSCLC (Ye et al., 2020). These can serve as potential diagnostic and therapeutic biomarkers for NSCLC. Moreover, the interactions among the identified DEGs can help better understand the underlying molecular mechanism of NSCLC. The identification of DEGs in NSCLC can contribute to the concept of personalised medicine.

In this study, bioinformatics analysis is performed to identify the significant cellular pathway correlated with NSCLC and its association with P16INK4a gene. We investigate the correlation between p16INK4a expression and genes involved in the ECM pathway in early-stage NSCLC.

## Methods and Protocols

### Data Collection

Three publicly available microarray datasets GSE19188, GSE27262, GSE118370 were considered for this study. All of them were established using the GPL570 [HG-U133_Plus_2] Affymetrix Human Genome U133 Plus 2.0. GSE19188 consisted of 91 lung cancer and 65 adjacent normal lung tissue, GSE27262 comprised 25 lung cancer and 25 adjacent paired normal lung tissue, and GSE118370 had 6 lung adenocarcinoma tissues and 6 paired normal lung tissues. Each of the datasets were individually categorised as test samples and controls and processed using GEO2R for identification of DEGs in NCBI. The selected datasets of all tumour samples and their control samples were analysed using GEO2R.

GEO2R is an R-based web tool that enables GEO users to compare multiple sets of samples from the queried GEO series in order to detect differentially expressed genes (DEGs) across the selected samples. The tool utilizes the GEO query and limma package from the Bioconductor project, which is widely recognized for its statistical analysis capabilities in identifying DEGs. DEGs were identified based on the cutoff criteria of P-value < 0.05 and |log fold change| > 2. To visualize the upregulated and downregulated genes, a volcano plot was created using the ggplot2 package in R. Analysed data was then downloaded into worksheets for further processing.

### Data Processing

The DEGs were further manually filtered using two criteria, P-value <0.05 and |log fold change| >2 for upregulated genes and |log fold change| <2 for downregulated genes. Volcano plots are constructed to represent the upregulated and downregulated genes observed using the ggplot2 package in R. After obtaining the DEGs through GEO2R for all three datasets, common DEGs are selected through a Venn diagram, using Venny software version 2.0.2, to identify the common DEGS in NSCLC between the three datasets. Upregulated and downregulated DEGs were also identified.

### GO And KEGG Analysis

To conduct gene ontology (GO) enrichment analysis of the shared 149 common DEGs from all three datasets were extracted and gene enrichment analysis was performed using Shiny go software, version 0.77. A significance threshold of <0.05 was applied to all the processes’ P-values. Shiny GO 0.77, developed by South Dakota State University, is a user-friendly bioinformatics tool for biologists. It aids in gene list analysis, identifying enriched Gene Ontology (GO) terms and pathways. The tool offers visualization options, integrates with iDEP and STRING-db, and is accompanied by a paper by (Ge et al., 2020) for reference.

### Protein-protein interactions (PPI) analysis

STRING is a database of known and predicted protein-protein interactions. The interactions include direct (physical) and indirect (functional) associations; they stem from computational prediction, from knowledge transfer between organisms, and from interactions aggregated from other (primary) databases. Parameters of a minimum interaction score of ≥ 0.4 were used when constructing the PPI network for differentially expressed genes (DEGs).

### Pathway Analysis

Reactome annotated data describe reactions possible if all annotated proteins and small molecules were present and active simultaneously in a cell. By overlaying an experimental dataset on these annotations, we performed a pathway over-representation analysis. By overlaying quantitative expression data or time series, we visualized the extent of change in affected pathways and its progression. A binomial test was used to calculate the probability shown for each result, and the p-values are corrected for the multiple testing (Benjamini–Hochberg procedure) that arises from evaluating the submitted list of identifiers against every pathway.

Reactome is a curated database of pathways and reactions in human biology. Reactions can be considered as pathway ‘steps’. Reactome defines a ‘reaction’ as any event in biology that changes the state of a biological molecule. Binding, activation, translocation, degradation and classical biochemical events involving a catalyst are all reactions. Orthologous reactions inferred from annotation for Homo sapiens are available for 14 non-human species including mouse, rat, chicken, puffer fish, worm, fly and yeast. Pathways are represented by simple diagrams following an SBGN-like format.

## Results

### Identification of DEGs

The three microarray datasets were processed individually for DEGs using GEO2R tool in NCBI. After filtration criteria, GSE19188 data contained a total of 635 DEGs out of which 465 are upregulated and 170 are downregulated. GSE27262 data contained a total of 671 DEGs out of which 487 are upregulated and 184 downregulated. GSE118370 data contained a total of 1,323 DEGs out of which 881 are upregulated and 442 downregulated.

Volcano plots were extracted from GEO2R results representing upregulated and downregulated DEGs as shown in Figure 2. Figure 2.A. represents DEGs from microarray dataset GSE19188, and Figure 2.B. represents DEGs from microarray dataset GSE27262, Figure 2.C. represents DEGs from microarray dataset and GSE118370.

**Figure 2.**
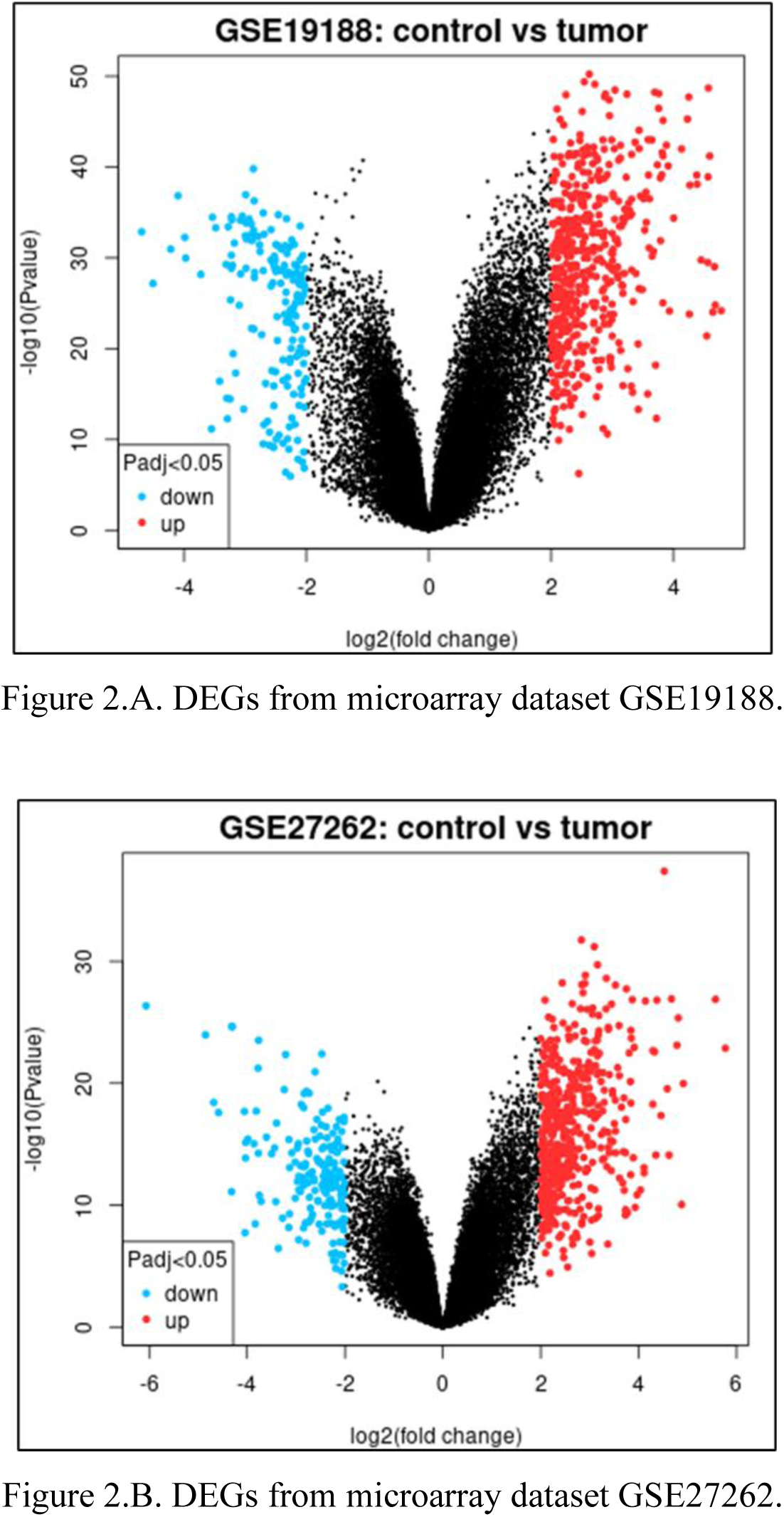
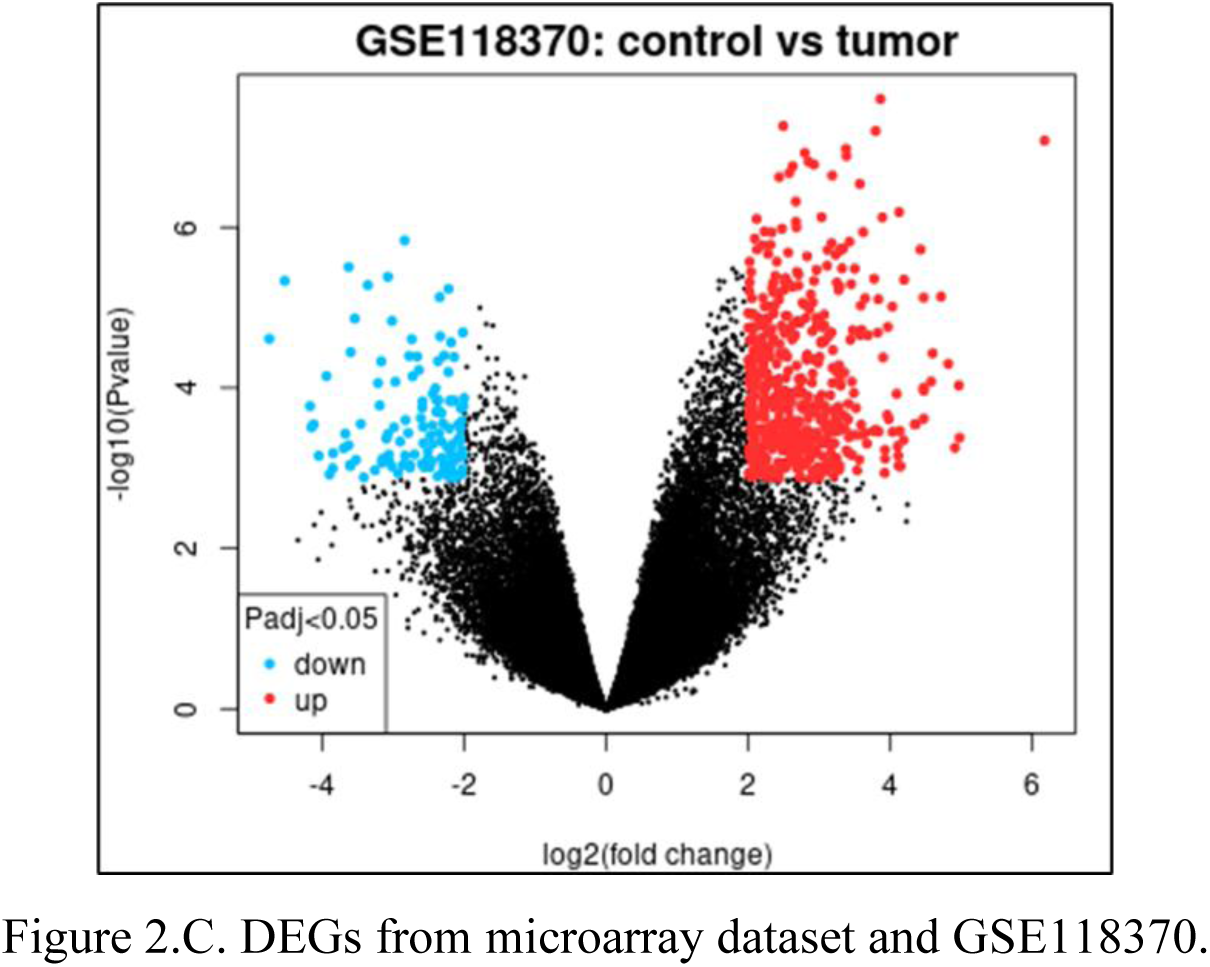
Volcano diagrams showcasing the significance of statistical data (-log10 P value) compared to the extent of alteration (log2 fold change, representing genes with differential expression. Notably, genes marked in red indicate upregulation, while those in blue represent downregulation, with a default adjusted p-value cutoff of 0.05. Figure 2.A. represents DEGs from microarray dataset GSE19188, Figure 2.B. represents DEGs from microarray dataset GSE27262, Figure 2.C. represents DEGs from microarray dataset and GSE118370.

Combining three data sets to identify common DEGs, Venn diagrams, shown in Figure 3.A, show that 149 common DEGs were present in “GSE19188 “, “GSE27262 “ and “GSE118370 “ combined, Venn diagrams are also used to the 22 common DEGs that were downregulated in “GSE19188 DR”, “GSE27262 DR” and “GSE118370 DR” as shown in Figure 3.B., Figure 3.C. represents Venn diagram to identify 127 common DEGs were upregulated in “GSE19188 UR”, “GSE27262 UR” and “GSE118370 UR”.

**Figure 3.**
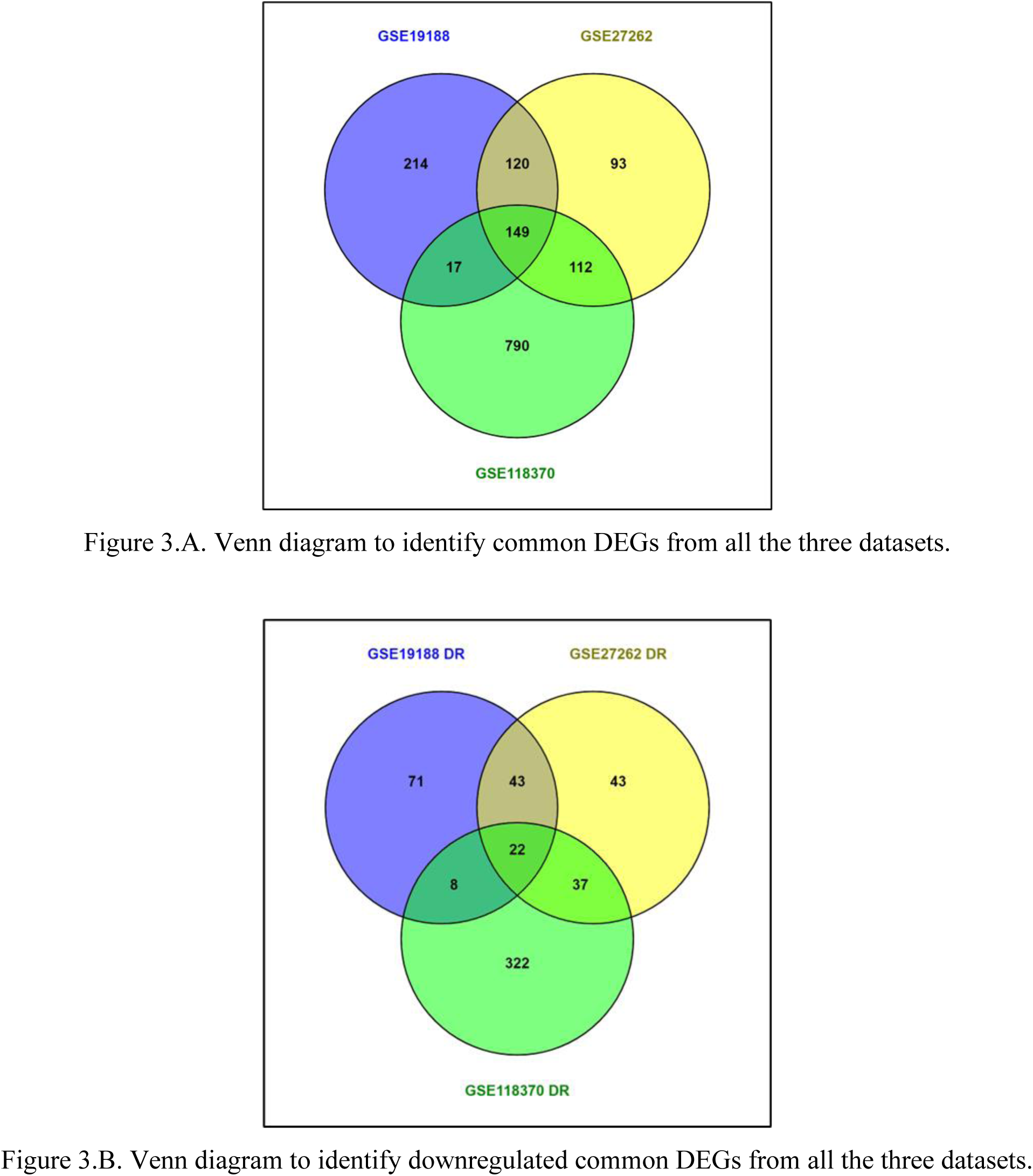
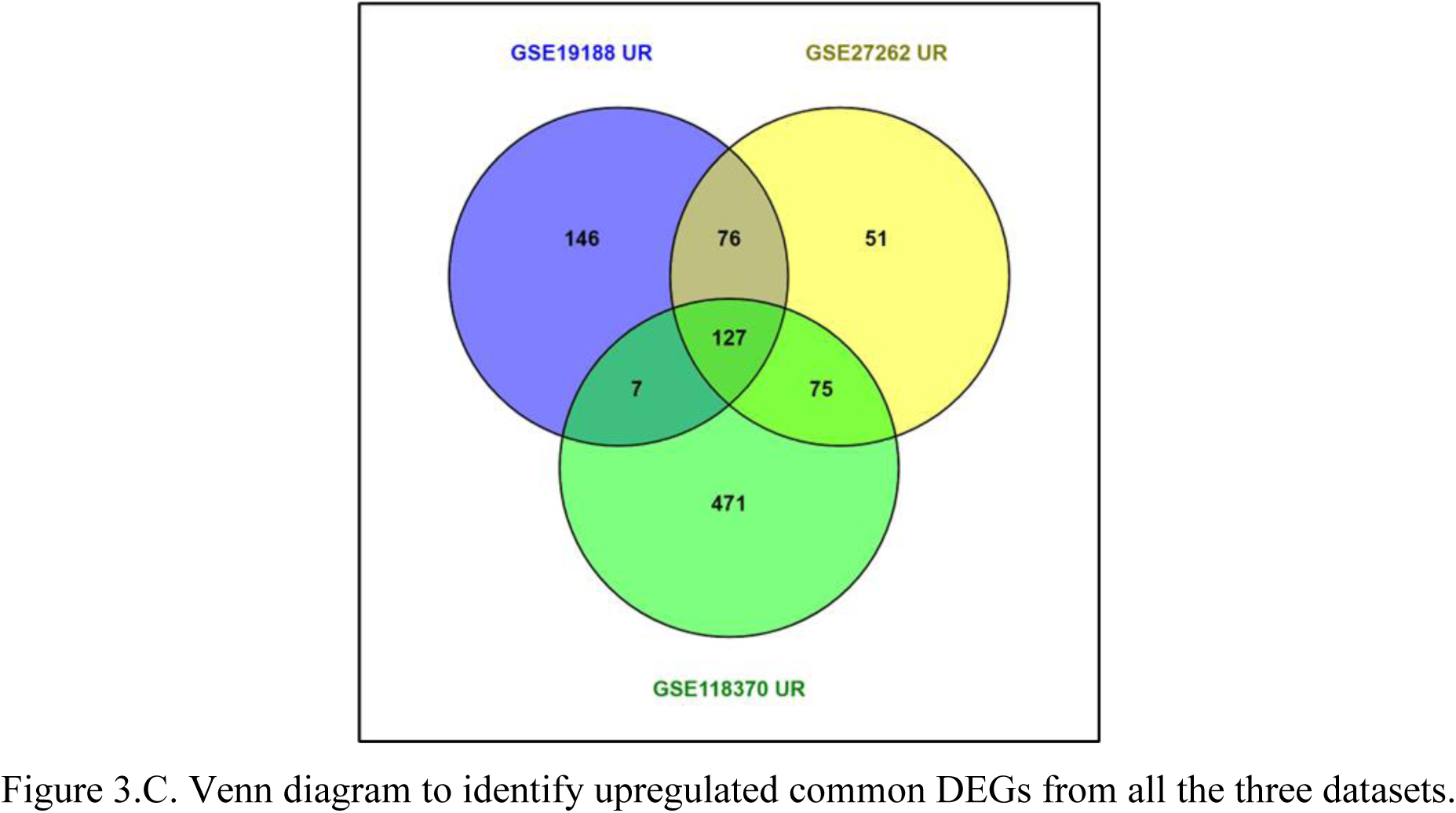
Venn diagram to identify common DEGs. Each circle represents DEGs from microarray datasets GSE19188 (violet), GSE27262 (yellow), and GSE118370 (green) respectively. The numbers in the intersection represents the common DEGs for all the datasets. Figure 3.A. represents Venn diagram to identify common DEGs from all the three datasets. Figure 3.B. represents Venn diagram to identify downregulated common DEGs from all the three datasets. Figure 3.C. represents Venn diagram to identify upregulated common DEGs from all the three datasets.

### Gene set analysis of DEGs

Gene Ontology analysis showed the common DEGs for biological processes were mainly enriched in ECM pathway. Figure 4. Pathway Enrichment analysis performed on 149 common DEGs with applied cutoff at p < 0.05. the X-axis represents the fold enrichment values.

**Figure 4.**
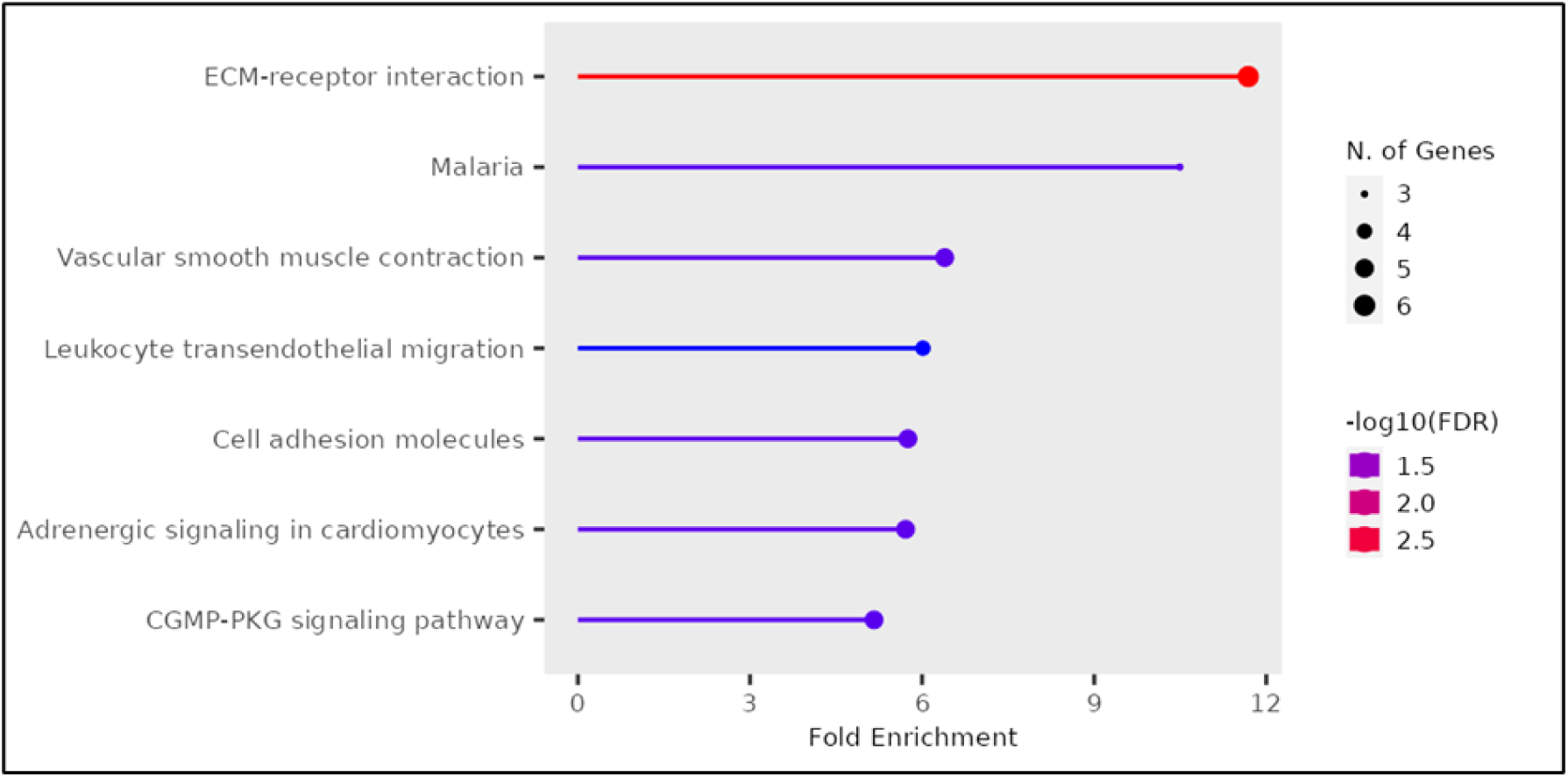
Pathway Enrichment analysis performed on 149 common DEGs with applied cutoff at p < 0.05. the X-axis represents the fold enrichment values. The Y-axis represents the top molecular pathways for common DEGs in descending order of the fold enrichment. The density of the tip represents the number of genes involved in respective pathways. The colour represents the significance.

### PPI network analysis

ECM receptor interaction - ECM receptors, particularly integrins, are involved in signalling pathways that influence cell behaviour. Dysregulation of integrin-mediated signalling can impact cell cycle progression, which may indirectly affect p16INK4a levels or activity. ECM also undergoes remodelling in response to various cues, and this process can affect tumour development and progression. Altered ECM composition may influence p16INK4a expression or function, indirectly impacting cell cycle regulation. ECM plays a critical role in creating the tumour microenvironment, which affects cancer cell behaviour and responses to therapy. Changes in the tumour microenvironment can influence p16INK4a expression and its tumour-suppressive effects.

### Pathway analysis

This analysis entails an overrepresentation assessment, which employs a statistical test based on the hypergeometric distribution. Its primary aim is to ascertain whether specific Reactome pathways exhibit a higher representation (enrichment) within the provided dataset. Essentially, it addresses the question of whether the list of proteins in the dataset demonstrates a more substantial presence in Pathway X than would be anticipated by random chance. The outcome of this test is a probability score, which is further adjusted for false discovery rate using the Benjamini-Hochberg method.

Among the sample’s 149 identifiers, 97 were identified within the Reactome database, and collectively, they interacted with 458 distinct pathways. Table 1 shows the 25 most relevant pathways sorted by p-value. All non-human identifiers have been converted to their human equivalent. Figure 5 Genome-wide overview of the pathway analysis. Reactome pathways are arranged in a hierarchy. The centre of each of the circular “bursts” is the root of one top-level pathway. Each step away from the centre represents the next level lower in the pathway hierarchy.

**Figure 5.**
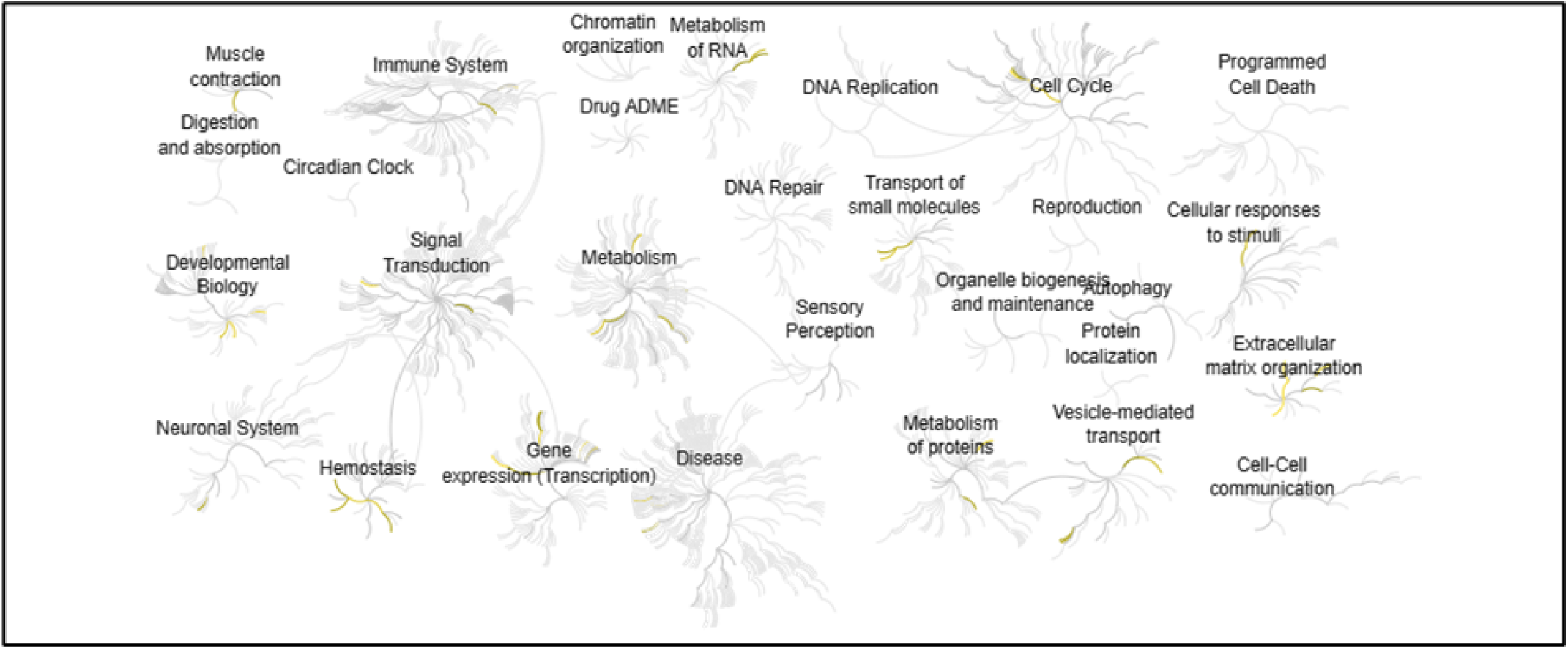
Genome-wide overview of the pathway analysis. Reactome pathways are arranged in a hierarchy. The centre of each of the circular “bursts” is the root of one top-level pathway. Each step away from the centre represents the next level lower in the pathway hierarchy. The colour code denotes over-representation of that pathway in our input dataset. Light grey signifies pathways which are not significantly over-represented.

**Table 1.** Shows the 25 most relevant pathways sorted by p-value. (* False Discovery Rate)

| SI<br>No. | Pathway name | Entities |  |  |  | Reactions |  |
| --- | --- | --- | --- | --- | --- | --- | --- |
|  |  | found | ratio | p-value | FDR* | found | ratio |
| 1 | Integrin cell surface interactions | 7/86 | 0.005<br>64600<br>8 | 7.86E-05 | 0.0375097<br>87 | 15/55 | 0.0038442<br>72 |
| 2 | Polo-like kinase mediated events | 4/23 | 0.001<br>50997<br>9 | 1.70E-04 | 0.040575 | 4/15 | 0.0010484<br>38 |
| 3 | FOXO-mediated transcription of cell cycle genes | 4/27 | 0.001<br>77258<br>4 | 3.12E-04 | 0.0496455<br>83 | 4/22 | 0.0015377<br>09 |
| 4 | Extracellular matrix organization | 12/328 | 0.021<br>53361<br>3 | 5.33E-04 | 0.0634460<br>61 | 85/319 | 0.0222967<br>78 |
| 5 | Interleukin-33 signalling | 2/4 | 2.63E<br>-04 | 0.0010534<br>46 | 0.1000773<br>71 | 2/2 | 1.40E-04 |
| 6 | Collagen degradation | 5/69 | 0.004<br>52993<br>7 | 0.0014040<br>46 | 0.1109196<br>34 | 14/34 | 0.0023764<br>59 |
| 7 | Degradation of the extracellular matrix | 7/148 | 0.009<br>71638<br>7 | 0.0019279<br>9 | 0.1311032<br>9 | 32/105 | 0.0073390<br>65 |
| 8 | Formation of lateral plate mesoderm | 2/8 | 5.25E<br>-04 | 0.0040866<br>22 | 0.2411106<br>78 | 4/5 | 3.49E-04 |
| 9 | RUNX3 Regulates Immune Response and Cell Migration | 2/10 | 6.57E-04 | 0.00628853 | 0.256744065 | 1/5 | 3.49E-04 |
| 10 | Muscle contraction | 8/232 | 0.015231092 | 0.006319307 | 0.256744065 | 14/53 | 0.00370448 |
| 11 | Interaction between L1 and Ankyrins | 3/33 | 0.002166492 | 0.007089649 | 0.256744065 | 2/4 | 2.80E-04 |
| 12 | TP53 Regulates Transcription of Cell Cycle Genes | 4/56 | 0.004267332 | 0.007452009 | 0.256744065 | 4/42 | 0.002935626 |
| 13 | Calcitonin-like ligand receptors | 2/11 | 7.22E-04 | 0.007551296 | 0.256744065 | 4/8 | 5.59E-04 |
| 14 | Negative regulation of activity of TFAP2 (AP-2) family transcription factors | 2/11 | 7.22E-04 | 0.007551296 | 0.256744065 | 3/6 | 4.19E-04 |
| 15 | Transcriptional regulation of white adipocyte differentiation | 5/109 | 0.007155987 | 0.009536586 | 0.26146099 | 4/19 | 0.001328021 |
| 16 | Defective B3GALTL causes PpS | 3/39 | 0.002560399 | 0.011124069 | 0.26146099 | 1/1 | 6.99E-05 |
| 17 | Cell surface interactions at the vascular wall | 8/257 | 0.016872374 | 0.011248716 | 0.26146099 | 28/65 | 0.004543231 |
| 18 | Defective pro-SFTPC<br>causes SMDP2 and<br>RDS | 1/1 | 6.57E<br>-05 | 0.0116182<br>88 | 0.2614609<br>9 | 1/1 | 6.99E-05 |
| 19 | O-glycosylation of<br>TSR domain-<br>containing proteins | 3/41 | 0.002<br>69170<br>2 | 0.0127086<br>9 | 0.2614609<br>9 | 2/2 | 1.40E-04 |
| 20 | G2/M Transition | 7/212 | 0.013<br>91806<br>7 | 0.0129102<br>63 | 0.2614609<br>9 | 34/78 | 0.0054518<br>77 |
| 21 | Mitotic G2-G2/M<br>phases | 7/214 | 0.014<br>04937 | 0.0135267<br>64 | 0.2614609<br>9 | 34/80 | 0.0055916<br>68 |
| 22 | Triglyceride<br>catabolism | 3/42 | 0.002<br>75735<br>3 | 0.0135469<br>02 | 0.2614609<br>9 | 6/17 | 0.0011882<br>3 |
| 23 | Formation of Fibrin<br>Clot (Clotting<br>Cascade) | 3/43 | 0.002<br>82300<br>4 | 0.0144159<br>26 | 0.2614609<br>9 | 4/61 | 0.0042636<br>47 |
| 24 | Collagen chain<br>trimerization | 3/44 | 0.002<br>88865<br>5 | 0.0153158<br>96 | 0.2614609<br>9 | 3/28 | 0.0019570<br>84 |
| 25 | FMO oxidises<br>nucleophiles | 2/16 | 0.001<br>05042 | 0.0153800<br>58 | 0.2614609<br>9 | 2/3 | 2.10E-04 |

#### 1. Integrin cell surface interactions

Highly significant was the integrin cell surface interactions as mentioned in the table and Figure 6 Integrin cell surface interactions diagram. The extracellular matrix (ECM) is a complex network of molecules providing mechanical support and influencing cell behaviour. It includes collagens, fibronectin, and laminins as key components, along with molecules like vitronectin and thrombospondin. Cell-ECM interactions are mediated by integrins, consisting of alpha and beta subunits forming heterodimers. In humans, 18 alpha and 8 beta subunits combine to create 24 receptor variants, categorized into beta1, beta2/beta7, and beta3/alphaV integrin families. These integrins play vital roles in cell adhesion, with beta1 binding to RGD, collagen, or laminin, beta2/beta7 in leukocyte interactions, and beta3/alphaV in RGD recognition, including aIIbb3 on platelets and others.

**Figure 6.**
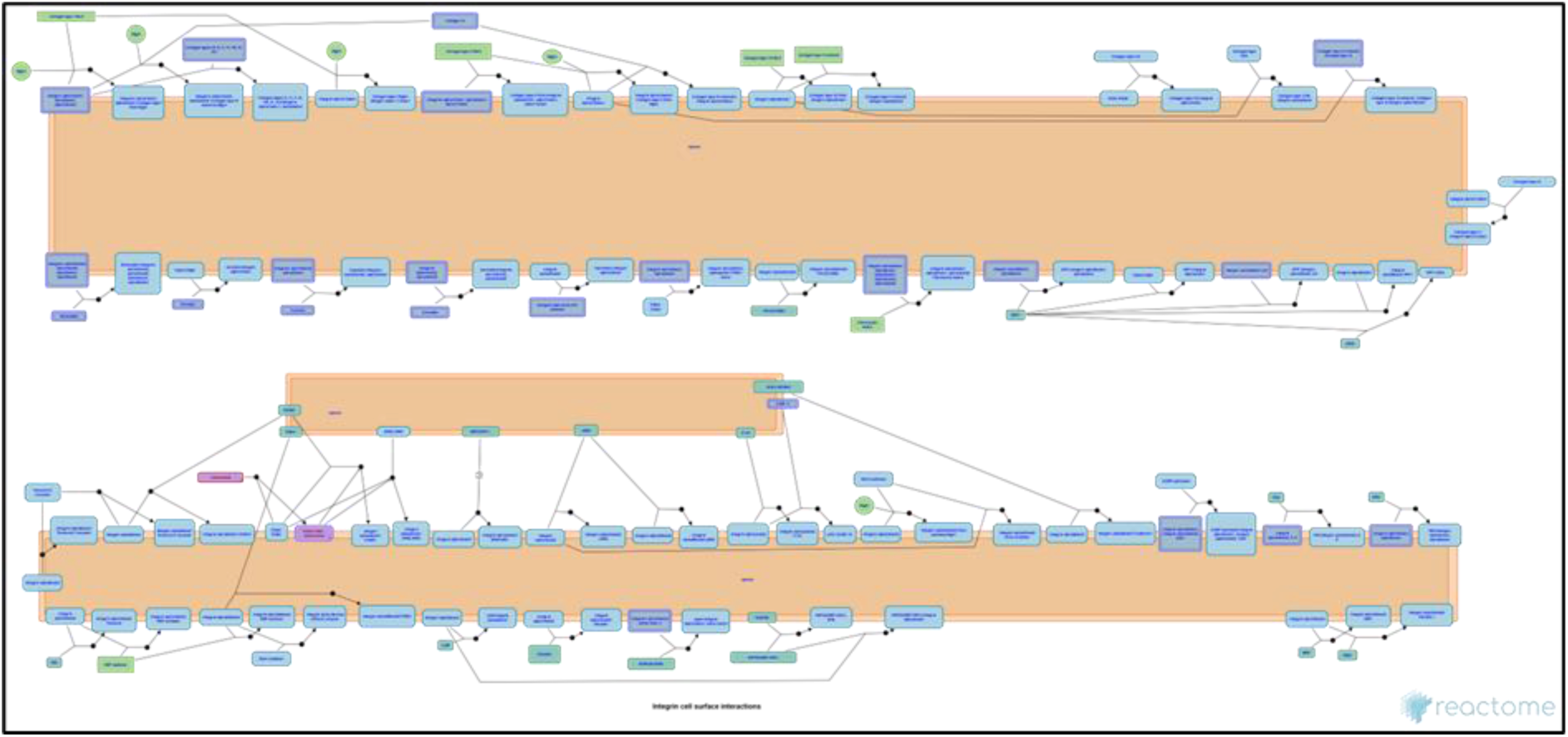
Integrin cell surface interactions diagram (*Reactome | Integrin Cell Surface Interactions*, n.d.). The extracellular matrix (ECM) is a diverse network of molecules providing strength and influencing cell behaviour. Integrins mediate cell-ECM interactions.

#### 2. Polo-like kinase mediated events

In the nucleoplasm, as shown in Figure 7 Polo-like kinase mediated events diagram. At the stage of entering the mitotic stage, Plk1 has dual roles in regulating key proteins. It phosphorylates and activates Cdc25C phosphatase while phosphorylating and reducing Wee1A activity. Plk1 also inhibits Myt1 through phosphorylation. Cyclin B1-bound Cdc2, which these proteins target, forms a feedback loop by phosphorylating them in return. This phosphorylation creates binding sites for Plk1’s polo-box domain, facilitating Plk1’s control over these proteins and ultimately activating Cdc2-Cyclin B1. Additionally, Plk1 phosphorylates and activates the transcription factor FOXM1, which promotes the expression of genes crucial for G2/M transition, including PLK1 itself, establishing a positive feedback loop.

**Figure 7.**
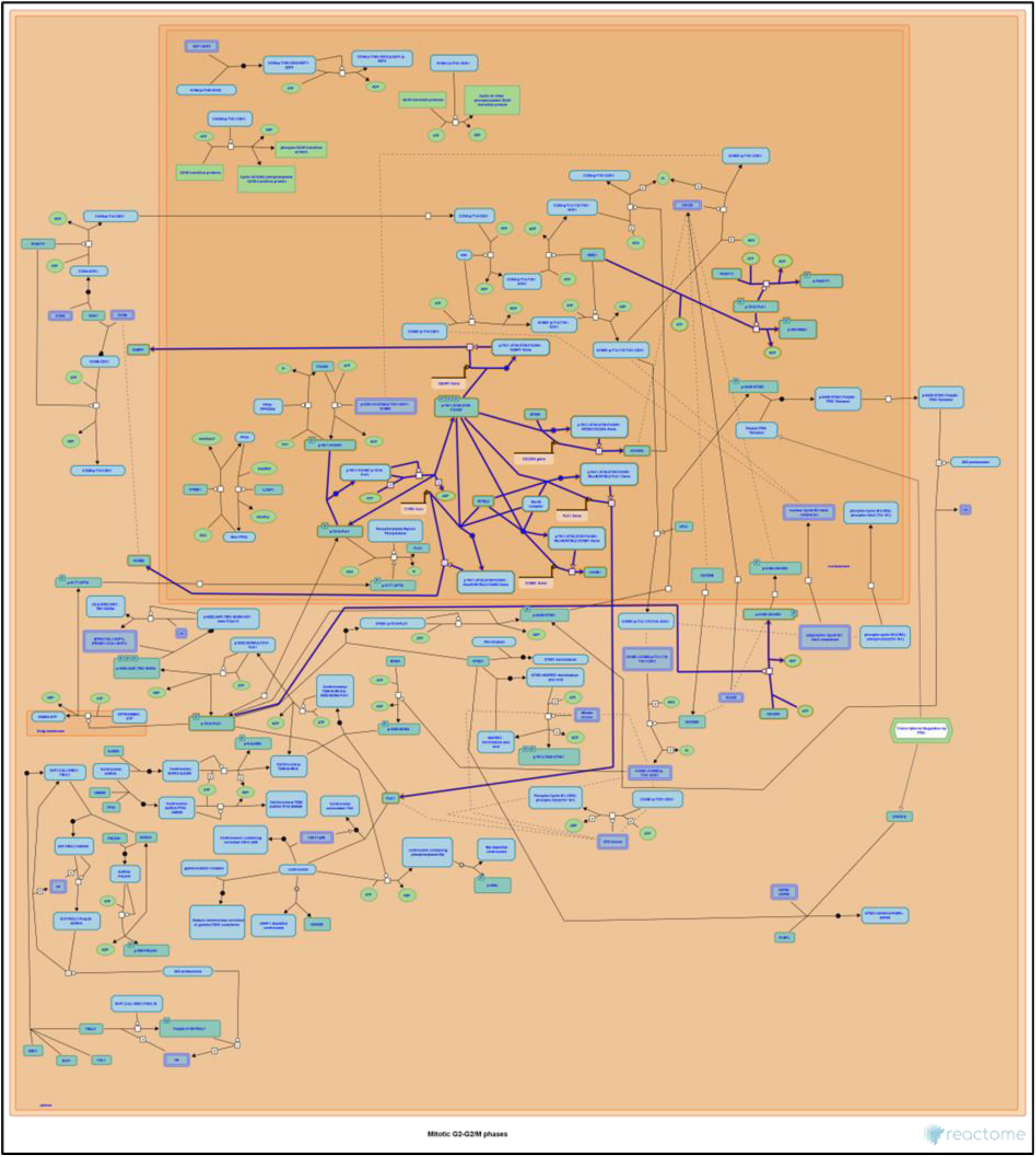
Polo-like kinase mediated events diagram. Plk1 orchestrates mitotic entry by phosphorylating Cdc25C, Wee1A, and Myt1, creating a feedback loop and activating essential cell cycle components.

#### 3. FOXO-mediated transcription of cell cycle genes

FOXO transcription factors play crucial roles in controlling cell proliferation as shown in Figure 8 FOXO-mediated transcription of cell cycle genes diagram.. They induce the expression of various genes that act as inhibitors of cell proliferation in different cell types. For instance, FOXOs stimulate the transcription of CDKN1A (p21Cip1) and CDKN1B (p27Kip1), both of which are cyclin-dependent kinase (CDK) inhibitors. They can also cooperate with the SMAD2/3:SMAD4 complex to induce CDKN1A transcription in response to TGF-beta signalling. Additionally, FOXOs directly stimulate the transcription of GADD45A, RBL2 (p130), CCNG2, BTG1, CAV1, KLF4, and MSTN genes, which are all involved in inhibiting cell proliferation and promoting quiescence or differentiation. FOXO transcription factors thus play a critical role in regulating cell growth and differentiation.

**Figure 8.**
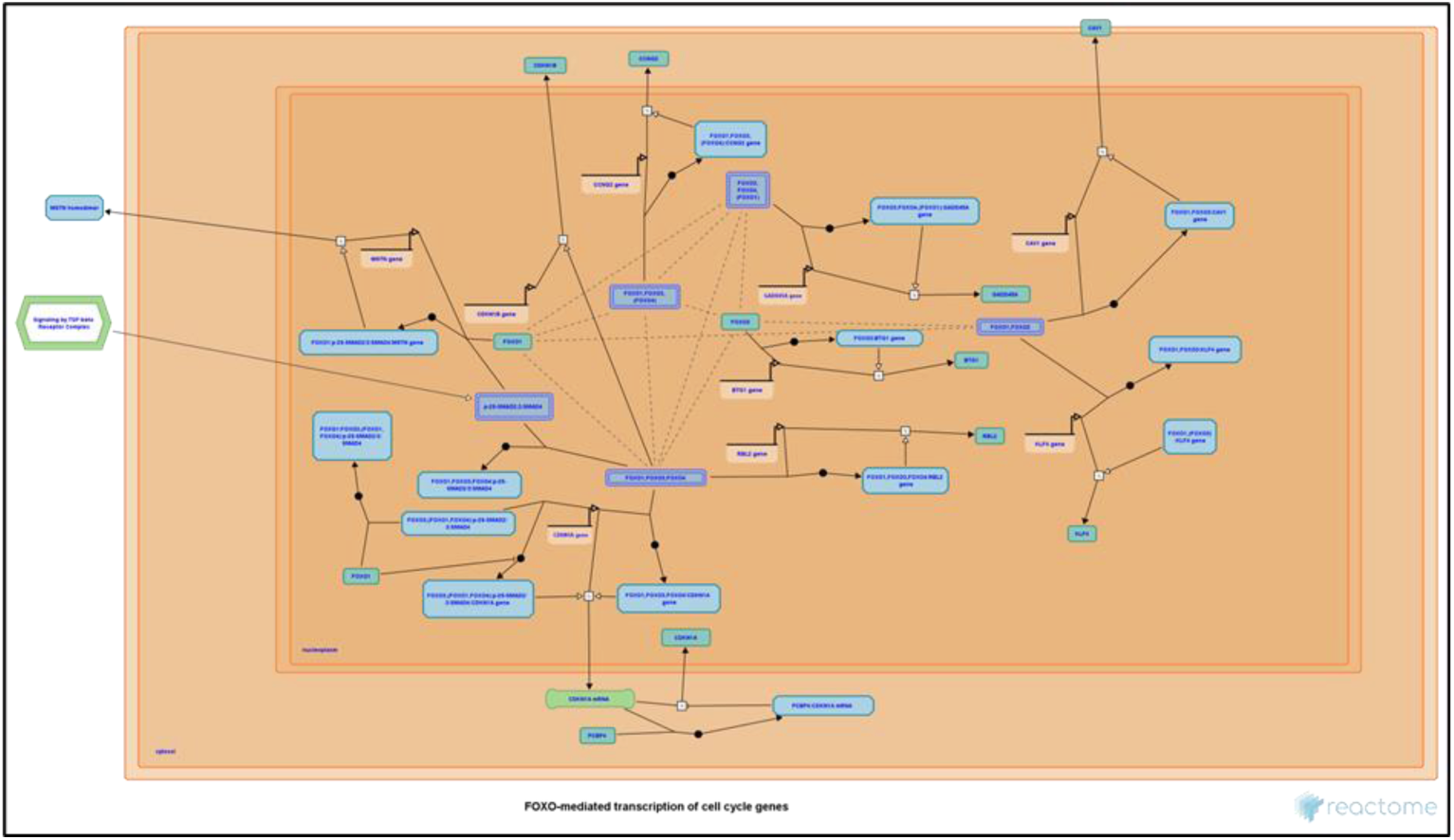
FOXO-mediated transcription of cell cycle genes diagram. FOXO transcription factors activate genes that inhibit cell proliferation in various cell types, including erythroid progenitors and neuroepithelial progenitor cells. They directly stimulate the transcription of CDK inhibitors like CDKN1A and CDKN1B, as well as genes like GADD45A, RBL2 (p130), CCNG2, BTG1, CAV1, KLF4, and MSTN involved in controlling cell growth and quiescence. These actions contribute to the regulation of cell proliferation and differentiation.

#### 4. Extracellular matrix organization

The extracellular matrix (ECM) as represented in Figure 9, is a critical component found in all mammalian tissues, composed mainly of fibrous proteins like collagen, elastin, fibronectin, and laminins, embedded within a gel-like structure formed by anionic proteoglycan polymers. Apart from its structural role, the ECM profoundly influences cell behaviours such as adhesion, migration, differentiation, and cell death. This dynamic composition is constantly remodelled and regulated by matrix metalloproteinases (MMPs) and growth factors. ECM remodelling plays a vital role in various processes, including stem cell niches, angiogenesis, and wound repair. Dysregulation of ECM dynamics can lead to cell proliferation issues, tissue fibrosis, and cancer. Collagen, elastin, fibrillin, and other ECM proteins contribute to tissue strength and flexibility, while proteoglycans, like aggrecan, help maintain cartilage’s turgid nature. MMPs, ADAMTS, and other enzymes control ECM remodelling, with TIMPs acting as potent MMP inhibitors.

**Figure 9.**
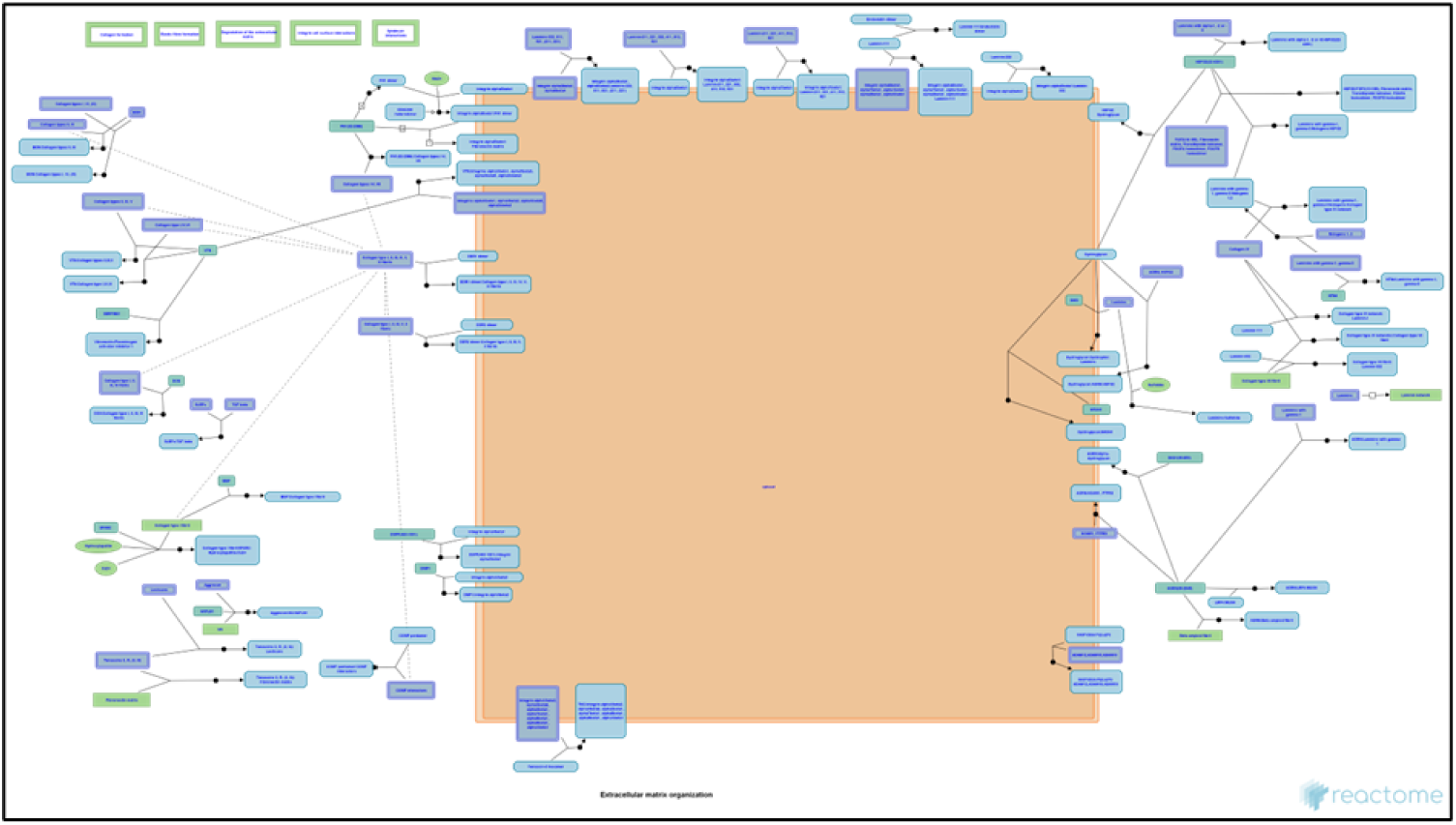
Extracellular matrix organization diagram. ECM, vital tissue framework, comprises fibrous proteins, proteoglycans, and enzymes.

#### 5. Interleukin-33 signalling

The Interleukin-33 signalling is represented in Figure 10 Interleukin-33 signalling diagram. Interleukin-33 (IL33) is an Interleukin-1 family cytokine and functions as an alarmin, released during cell damage, serving as an endogenous danger signal. Full-length IL33 can be biologically active and undergoes increased potency, up to 30-fold, when cleaved at the N-terminus by inflammatory proteases like Cathepsin G (CTSG) and Neutrophil elastase (ELANE). However, some argue that processing may inactivate IL33. IL33 has a nuclear localization sequence, allowing it to translocate to the nucleus where it binds to heterochromatin. IL33’s binding to its receptor Interleukin-1 receptor-like 1 (IL1RL1 or ST2) initiates various cellular signalling pathways. It is primarily released into the extracellular space during cell injury or death, rather than actively secreted. Soluble IL1RL1 (ST2V) shares some components with IL1RL1 but lacks its transmembrane and intracellular parts. The IL33-IL1RL1 complex recruits a co-receptor, typically IL1 receptor accessory protein (IL1RAP or IL-1RAcP).

**Figure 10.**
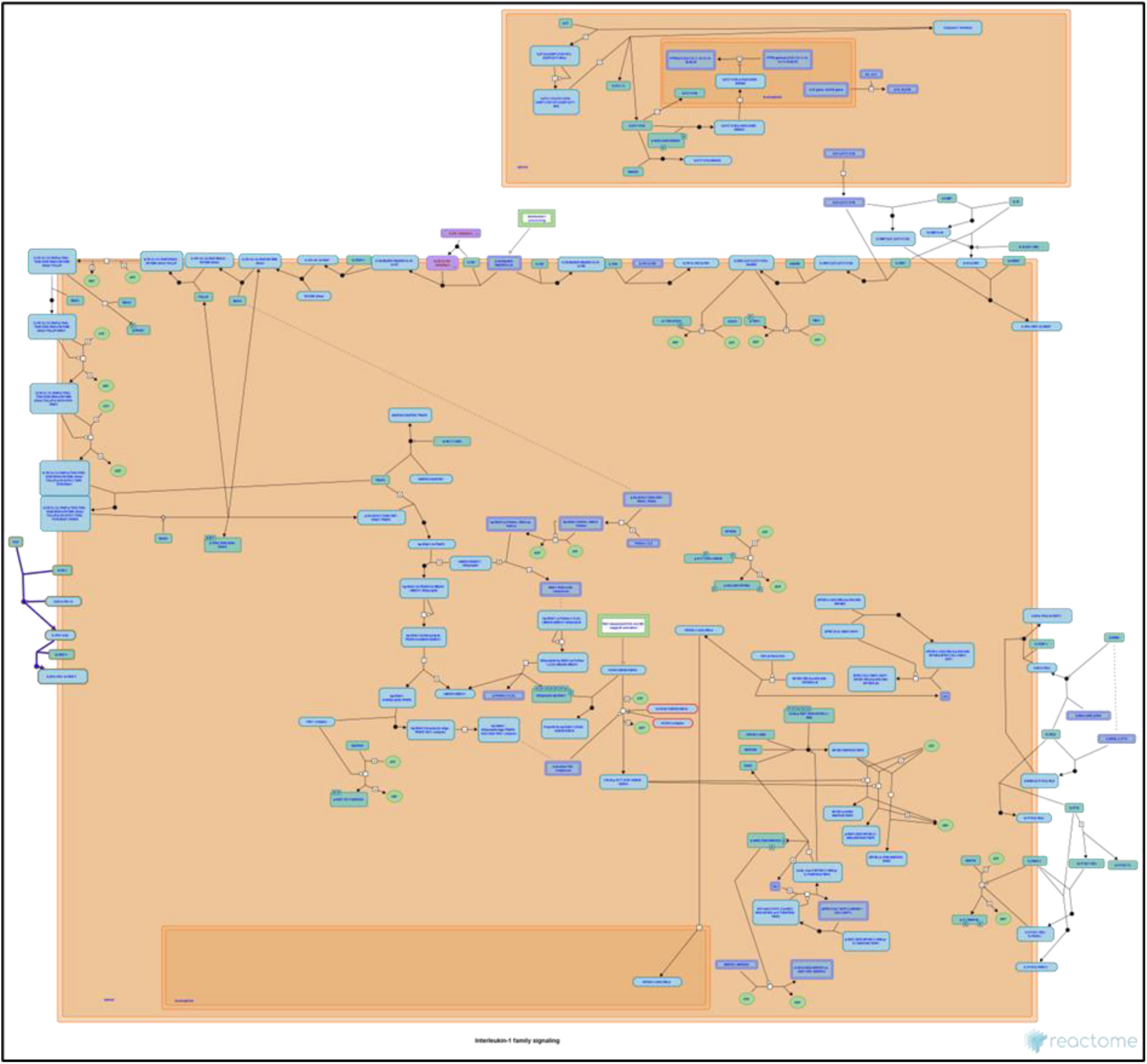
Interleukin-33 signalling diagram. Interleukin-33 (IL33) functions as an alarmin, released during cell damage, and engages cellular signalling through IL1RL1 (ST2) and IL1RAP.

#### 6. Collagen degradation

Collagen fibril diameter and arrangement vary with species, tissue type, and developmental stage. The degradation pathway is depicted in Figure 11. Collagen fibril lengths, particularly in tendons, can measure millimetres. These fibrils consist of around 350 collagen molecules in cross-section, narrowing to three molecules at the growth tip. Traditional collagenase action involves actively unwinding the triple helix (molecular tectonics) before cleaving the alpha2 chain and other chains. Recent research suggests a dynamic equilibrium between protected and vulnerable collagen states, where local unfolding allows collagenolysis by matrix metalloproteinases (MMPs). Collagen fibrils with cut chains become unstable, leading to gelatin formation. MMPs like MMP1, MMP8, and MMP13 play a significant role in collagen degradation, with specific cleavage sites on collagen chains. MMP14, a membrane-associated MMP, can cleave multiple collagen types.

**Figure 11.**
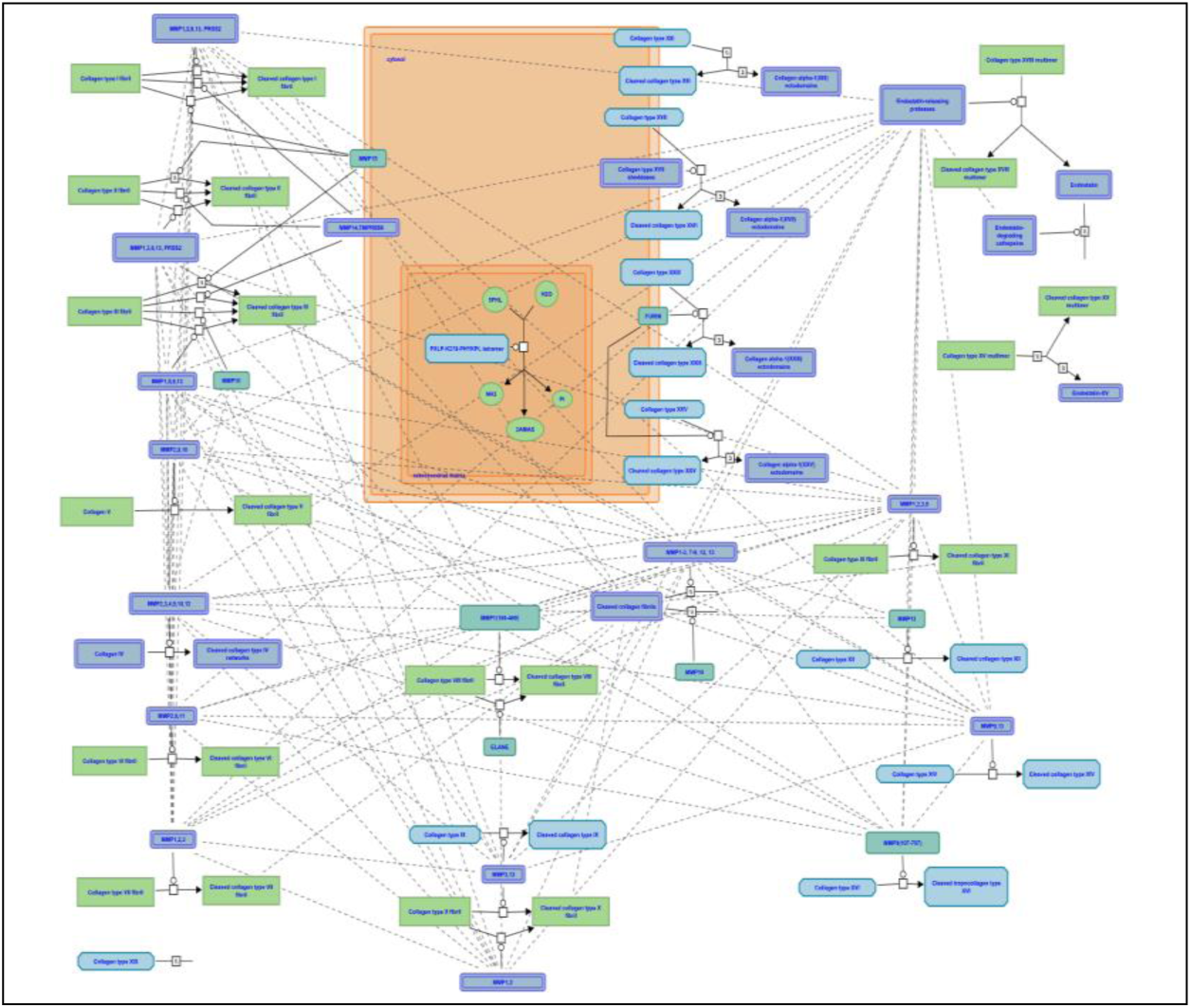
Collagen degradation pathway. Collagen fibril size varies by species and tissue type. Collagenase enzymes like MMP1, MMP8, and MMP13, also known as collagenases I, II, and III, initiate collagen breakdown. They cleave collagen alpha chains at specific Gly-Ile/Leu sites, primarily in collagens I, II, and III, influencing tissue remodelling processes. MMP14, a membrane-associated MMP, can also cleave these collagens.

## Discussion

This study delves into the crucial issue of non-small cell lung cancer (NSCLC), a significant global health concern with high mortality rates and often late-stage diagnoses. Despite medical advances, the prognosis for NSCLC remains suboptimal, emphasizing the urgency of improving early diagnosis and treatment strategies.

The research focuses on P16INK4a, a tumour suppressor gene implicated in NSCLC. Understanding its role and the molecular mechanisms associated with it could potentially pave the way for targeted therapies and improved patient outcomes. The use of bioinformatics tools to identify differentially expressed genes (DEGs) associated with NSCLC across multiple datasets is a commendable approach. This allows for the identification of potential biomarkers for early detection and personalized treatment options. The study’s efforts to explore gene ontology (GO) and Kyoto Encyclopaedia of Genes and Genomes (KEGG) pathway enrichment analyses further enhance our understanding of NSCLC’s underlying biological processes, with a notable emphasis on the extracellular matrix (ECM) pathway. Figure 9 shows the diagrammatic representation of extracellular matrix organization.

One of the key findings of this study is the identification of 149 common DEGs across multiple datasets, exploring protein-protein interactions (PPI) networks and conducting further validation experiments has identified that these DEGs have a significant role in the ECM pathway in NSCLC patients. To better understand the potential corelation between ECM pathway role of P16INK4a we conducted intensive literature review.

P16INK4a’s role in cell cycle regulation and tumour suppression can influence how cells interact with the ECM. Understanding the connections between p16INK4a and the ECM pathway is important for gaining insights into cancer development, tissue homeostasis, and potential therapeutic interventions. The p16INK4a gene is related to the ECM pathway through its involvement in cell cycle regulation and cell proliferation ECM receptors. Particularly, integrins are involved in signalling pathways that influence cell behaviour (Alberts et al., 2002). Dysregulation of integrin-mediated signalling can impact cell cycle progression, which may indirectly affect p16INK4a levels or activity.

The ECM also undergoes remodelling in response to various cues, and this process can affect tumour development and progression. Altered ECM composition may influence p16INK4a expression or function, indirectly impacting cell cycle regulation. The ECM plays a critical role in creating the tumour microenvironment, which affects cancer cell behaviour and responses to therapy (Brassart-Pasco et al., 2020). Changes in the tumour microenvironment can influence p16INK4a expression and its tumour-suppressive effects.

The CIP/KIP family is made up of three proteins: p21cip1/waf1, P27kip1, p57kip2 (Zohny et al., 2019). Sequence homology at the N-terminal domain which allows them to bind to both the cyclin and CDK. The INK 4 family inhibitors which include p15, p16, p18, and p19 are specific to CDK4 and its close isoform CDK6 and can bind to either the isolated CDK subunit or its complex with cyclin D (Cánepa et al., 2007). This is an exciting avenue that warrants more extensive investigation and experimentation. The development of targeted therapies holds promise for improving patient care and survival rates.

### Integrin cell surface interactions

Integrins are the receptors that mediate cell adhesion to ECM. Integrins consists of one alpha and one beta subunit forming a noncovalently bound heterodimer (Danen, 2013). 18 alpha and 8 beta subunits have been identified in humans that combine to form 24 different receptors. Figure 6 shows the diagrammatic representation of integrin cell surface interactions. The extracellular matrix (ECM) is a diverse network of molecules providing strength and influencing cell behaviour. Integrins mediate cell-ECM interactions.

The integrin dimers can be broadly divided into three families consisting of the beta1, beta2/beta7, and beta3/alphaV integrins (Geiger & Horwitz, 2008). beta1 associates with 12 alpha-subunits and can be further divided into RGD-, collagen-, or laminin binding and the related alpha4/alpha9 integrins that recognise both matrix and vascular ligands. beta2/beta7 integrins are restricted to leukocytes and mediate cell-cell rather than cell-matrix interactions, although some recognize fibrinogen. The beta3/alphaV family members are all RGD receptors and comprise aIIbb3, an important receptor on platelets, and the remaining b-subunits, which all associate with alphaV. It is the collagen receptors and leukocyte specific integrins that contain alpha A-domains. The Integrin cell surface interactions and P16INK4a are connected through their involvement in the influencing the ECM pathway.

### Polo-like kinase mediated events

At mitotic entry, Plk1 phosphorylates and activates Cdc25C phosphatase, whereas it phosphorylates and down-regulates Wee1A (Watanabe et al., 2004). Plk1 also phosphorylates and inhibits Myt1 activity (Inoue & Sagata, 2005). Cyclin B1-bound Cdc2, which is the target of Cdc25C, Wee1A, and Myt1, functions in a feedback loop and phosphorylates the latter components (Cdc25C, Wee1A, Myt1). The Cdc2-dependent phosphorylation provides docking sites for the polo-box domain of Plk1, thus promoting the Plk1-dependent regulation of these components and, as a result, activation of Cdc2-Cyclin B1. Figure 7 shows the diagrammatic representation of polo-like kinase mediated events.

PLK1 phosphorylates and activates the transcription factor FOXM1 which stimulates the expression of a number of genes needed for G2/M transition, including PLK1, thereby creating a positive feedback loop.

### FOXO-mediated transcription of cell cycle genes

FOXO transcription factors induce expression of several genes that negatively regulate proliferation of different cell types, such as erythroid progenitors (Bakker et al., 2004), (Wang et al., 2015) and neuroepithelial progenitor cells in the telencephalon (Seoane et al., 2004). Figure 8 shows a diagrammatic representation of FOXO-mediated transcription of cell cycle genes. Transcription of cyclin-dependent kinase (CDK) inhibitors CDKN1A (p21Cip1) is directly stimulated by FOXO1, FOXO3 and FOXO4. FOXO transcription factors can cooperate with the SMAD2/3: SMAD4 complex to induce CDKN1A transcription in response to TGF-beta signalling. FOXO transcription factors FOXO1, FOXO3 and FOXO4 stimulate transcription of the CDKN1B (p27Kip1) gene, but direct binding of FOXOs to the CDKN1B gene locus has not been demonstrated. Transcription of the retinoblastoma family protein RBL2 (p130), involved in the maintenance of quiescent (G0) state, is directly stimulated by FOXO1, FOXO3 and FOXO4. Transcription of the anti-proliferative protein CCNG2 is directly stimulated by FOXO1 and FOXO3, and possibly FOXO4.

### Interleukin-33 signalling

Interleukin-33 (IL33) cytokine is a member of the Interleukin-1 family. It can be classified as an alarmin because it is released into the extracellular space during cell damage. It acts as an endogenous danger signal (Liew et al., 2010). Figure 10 shows the diagrammatic representation of interleukin-33 signalling.

The gene product is biologically active (full-length IL33). Its potency has been reported to increase significantly (up to 30x) after cleavage at the N-terminus by inflammatory proteases such as Cathepsin G (CTSG) and Neutrophil elastase (ELANE) but others have suggested that processing inactivates IL33. IL33 can act as an extracellular ligand and an intracellular signalling molecule. Cell injury or death are the dominant mechanisms by which IL33 reaches the extracellular environment, IL33 is not actively secreted by cells. Because IL33 is expressed constitutively by endothelial and epithelial cells it is immediately available to the extracellular microenvironment after cell injury and necrosis. Increases in extracellular ATP or mechanical stress correlate with increased IL33 secretion by mast cells or cardiomyocytes, respectively. Soluble IL1RL1 (IL1RL1 Isoform C, ST2V) shares the extracellular components of IL1RL1, including the ligand binding domain, but lacks the transmembrane and intracellular components of IL1RL1. The IL33-IL1RL1 complex recruits a co-receptor, most commonly IL1 receptor accessory protein (IL1RAP, IL-1RAcP).

While the study’s findings are promising, it’s important to acknowledge some limitations. The research primarily relies on bioinformatics analyses of existing datasets. Experimental validation of the identified DEGs and their roles in NSCLC is necessary to confirm their clinical relevance fully. Additionally, the impact of genetic and environmental factors on NSCLC development and progression should be considered in future studies.

In conclusion, this study represents a significant step in the quest to improve the prognosis and treatment of NSCLC. It sheds light on the potential significance of P16INK4a and offers valuable insights into DEGs associated with NSCLC. To translate these findings into clinical practice, further research, including experimental validation and clinical trials, is needed. Ultimately, the study’s findings have the potential to make a substantial impact on global health by advancing early diagnosis and tailored treatment strategies for NSCLC patients.

## Acknowledgements

The authors would like to thank Dr. Monika Chauhan, Quick IsCool for review of the manuscript.

